# Comparing the appetitive learning performance of six European honeybee subspecies in a common apiary

**DOI:** 10.1101/2021.07.14.452344

**Authors:** R Scheiner, K Lim, MD Meixner, MS Gabel

## Abstract

The Western honeybee (*Apis mellifera* L.) is one of the most widespread insects with numerous subspecies in its native range. In how far adaptation to local habitats has affected the cognitive skills of the different subspecies is an intriguing question which we investigate in this study. Naturally mated queens of the following five subspecies from different parts of Europe were transferred to Southern Germany: *A. m. iberiensis* from Portugal, *A. m. mellifera* from Belgium, *A. m. macedonica* from Greece, *A*.*m. ligustica* from Italy and *A. m. ruttneri* from Malta. We also included the local subspecies *A*.*m. carnica* in our study. New colonies were built up in a common apiary where the respective queens were introduced. Worker offspring from the different subspecies was compared in classical olfactory learning performance using the proboscis extension response. Prior to conditioning we measured individual sucrose responsiveness to investigate whether possible differences in learning performances were due to a differential responsiveness to the sugar water reward. Most subspecies did not differ in their appetitive learning performance. However, foragers of the Iberian honeybee, *A. m. iberiensis*, performed significantly more poorly, despite having a similar sucrose responsiveness. We discuss possible causes for the low cognitive performance of the Iberian honeybees, which may have been shaped by adaptation to local habitat.

**Summary statement:** This study is the first to compare the associative learning performance of six honeybee subspecies from different European regions in a common apiary.

## Introduction

The natural range of the Western honeybee (*A. mellifera*) expands throughout Africa, Europe, Western and Central Asia to Western China in the East (Ruttner et al., 1978; Ruttner, 1988; Meixner et al., 2010; Sheppard and Meixner, 2003; Uzunov et al., 2015a,b; Chen et al., 2016). The intraspecific diversity of *Apis mellifera* is remarkable, with currently about 30 described subspecies (Ruttner, 1988; Cakmak et al., 2010; Bouga et al., 2011; Han et al., 2012; Chen et al., 2016). The different subspecies display diverse adaptations to a wide variety of geographic areas and environmental factors.

They can be grouped into five evolutionary lineages (A, M, C, O, and Y), based on morphometric and molecular studies (Ruttner, 1988; Moritz et al., 2007; Bouga et al., 2011; Han et al., 2012). Whereas the subspecies of lineage A are spread across Africa, those belonging to lineage Y originate in North-eastern Africa. Subspecies of lineage M are distributed widely in Western and Northern Europe, while lineage C comprises subspecies originating from South-Eastern Europe. Lineage O stems from the Near and Middle East. However, the native distribution of the different subspecies has been gravely altered by human interference and the current situation no longer represents the original distribution (Meixner et al., 2007; Meixner et al., 2010; Bouga et al., 2011). The worldwide demand for highly profitable honeybee colonies and the breeding efforts focusing on certain behavioral traits such as low aggressiveness have promoted the introduction of subspecies to locations outside their natural range, causing accidental and deliberate hybridization and leading to the jeopardization of numerous native populations of *A. mellifera* subspecies.

Selection has not only shaped the morphology of the different subspecies but also behavioral traits such as low aggressiveness and annual colony development cycles such as early spring development in the economically popular *A*.*m. carnica* (Adam, 1983; Ruttner, 1988; De La Rúa et al., 2009; Meixner et al., 2010; Uzunov et al., 2015a,b; Zammit-Mangion et al., 2017). However, huge behavioral differences have not only been observed between different subspecies of honeybees, but also among members of the same colony. A good example is individual responsiveness to sucrose, which has frequently been employed as a general indicator of the physiological state of a honeybee (Scheiner et al., 2004; Scheiner and Erber, 2009; Scheiner et al., 2013). It differs grossly between individuals performing different social tasks (Scheiner et al., 1999, 2001, 2017a,b; Reim and Scheiner, 2014; and between seasons (Scheiner et al., 2003). Importantly, it allows us to make predictions about the appetitive learning performance of the individual, because it correlates positively with cognitive performance in appetitive associative learning (Scheiner et al., 1999, 2001, 2003, 2005; Behrends and Scheiner, 2009). The more responsive a honeybee is to sucrose, the higher is her learning score, i.e. the better is her cognitive performance. So far, all of these experiments have been performed with workers of two subspecies of the Western honeybee, i.e. *A*.*m. carnica* and *A*.*m. ligustica*. Whether this relationship between cognitive performance and sensory responsiveness to sucrose is universal for different honeybee subspecies is an intriguing question. Similarly, it is an exciting open question whether different honeybee subspecies share similar cognitive capacities.

We here studied for the first time the appetitive olfactory learning performance in different subspecies of *Apis mellifera* maintained in a common garden apiary. We performed the experiments with colonies of six different European subspecies of *A. mellifera*, covering three of the five known evolutionary lineages: *A. m. mellifera* and *A. m. iberiensis* of lineage M, *A. m. carnica, A. m. macedonica* and *A. m. ligustica* from lineage C, and *A. m. ruttneri* from lineage A (Fig. 1). Our hypothesis was that the different subspecies should only differ in their appetitive learning performance if they differed in their responsiveness to sucrose, because this factor might be shaped by local adaptation to climate and it is an important determinant of learning performance (Scheiner et al. 1999, 2001a,b, 2005, 2013). Secondly, we aimed to investigate whether the correlation between individual sucrose responsiveness and appetitive learning performance described for *A*.*m. carnica* and *A*.*m. ligustica* would also be present in the other subspecies, pointing towards a general rule for appetitive associative learning.

**Figure 1.**
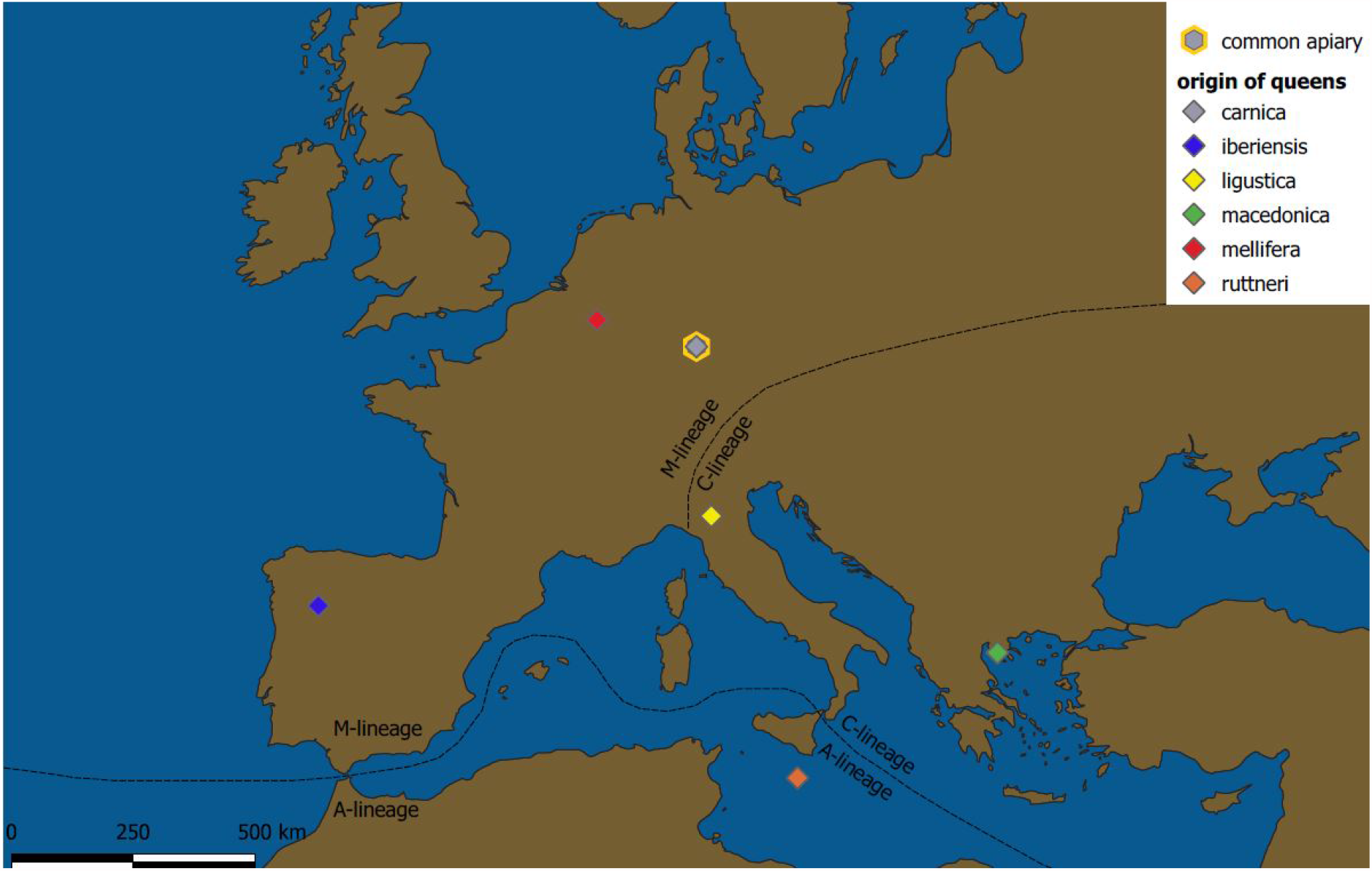
Map of lineage distribution ranges and points of origin of the different honeybee subspecies studied.

As a representative of lineage C, the Carniolan bee *A. m. carnica* (Fig. 1) is native to South-Eastern Austria and the North-Western Balkan Peninsula. It is nowadays one of the most commonly used subspecies in commercial beekeeping and bee science worldwide. From the late 19th century on, hives have been exported frequently from the Westernmost tip of its native distribution range, i.e. the eponymous Carniolan Alps, to various destinations (Ruttner, 1988). Due to its gentle temperament and low swarming tendency, characteristics regarded suitable for beekeeping, the “grey Carniolan bee” quickly became popular in Germany. Together with *A. m. ligustica* and the commercial hybrid Buckfast these bees have now by far the widest distribution range, often threatening other subspecies.

The *A. m. iberiensis* bees used in our experiment come from Bragança in Northern Portugal (Fig. 1). The natural distribution of this subspecies covers the whole Iberian Peninsula and the Balearic Islands (Ruttner, 1988; De La Rúa et al., 2001; Radloff et al., 2001; Zammit-Mangion et al., 2017), where it nowadays partially overlaps with imported commercial stock of other subspecies (Meixner et al., 2010). Ruttner (1988) describes different ecotypes adapted to both cold and warm climate conditions. Brood rearing is highly economical without wasting of resources and a good overall potential for applied beekeeping. However, this subspecies has also been described by ample use of propolis, a very nervous behavior on the combs and ferocious defensive behavior (Adam, 1983; Ruttner, 1988).

The original distribution of *A. m. mellifera* (lineage M) extended throughout central Europe North of the Alps including the United Kingdom, Ireland and Scandinavia in the North, all over France in the West and across Poland to the Ural mountain range in the East (Ruttner, 1988). Today, it has been replaced by other subspecies in large parts of its former distribution range (Jensen et al., 2005; De La Rúa et al., 2009; Meixner et al., 2010; Van Engelsdorp and Meixner, 2010; Ruottinen et al., 2014). This includes wide parts of Germany, where Carniolan stock (*A*.*m. carnica*) is now predominant. Various ecotypes have been described, which share a brood rhythm adapted to nectar flow, ample use of propolis and a nervous behavior on the combs with a tendency to defensiveness (Adam, 1983; Ruttner, 1988). Our *A. m. mellifera* bees stem from a breeding Apiary in Belgium (Fig. 1).

The subspecies *Apis mellifera ruttneri* (lineage A) is endemic to the archipelago of Malta, which constitutes its former and actual range of distribution (Fig. 1; Sheppard et al., 1997; Zammit-Mangion et al., 2017). Due to its small area of distribution and frequent imports of other commercially used subspecies, *A. m. ruttneri* is highly prone to genetic introgression (Sheppard et al., 1997; Zammit-Mangion et al., 2017). Its brood cycle is adjusted to seasonal nectar flows and xeric conditions, while colonies show a moderate use of propolis and sometimes fierce defensive behavior (Sheppard et al., 1997). While the distinct defensive behavior, especially under hot and dry weather conditions, is an unfavorable trait for their use in apiculture, *A. m. ruttneri* is also able to cope with predatory wasps and the challenging seasons of the respective habitat (Sheppard et al., 1997).

The natural distribution range of *A. m. macedonica* (lineage C) extends from Northern Greece across North Macedonia, Bulgaria, Romania and Moldova to Southern Ukraine in the North (Ruttner, 1988; Uzunov et al., 2015b). Ruttner (1988) describes *A. m. macedonica* as gentle, but sometimes inclined to swarm and susceptible to nosemosis. He also reports ample use of propolis and brood reduction as a reaction to unsuitable (especially hot and dry) weather conditions. Our bees derived from northern Greece (Fig. 1).

The original distribution range of *A. m. ligustica* (lineage C) covers the Apennine Peninsula (Ruttner, 1988). In recent times, however, *A. m. ligustica* has been spread around the globe for its beekeeping value. Apart from *A. m. carnica*, this subspecies is by far the most commonly used in commercial apiculture (De La Rúa et al., 2009; Meixner et al., 2015). Beekeepers appreciate its high fertility, gentleness, little usage of propolis and high productivity under good foraging conditions (Adam, 1983; Ruttner, 1988). Under poor foraging conditions or in cold climates, however, colonies of *A. m. ligustica* do not adjust their brood rearing: a trait resulting in lower honey yields for beekeepers or starvation of colonies through accelerated store consumption (Adam, 1983; Ruttner, 1988). Our *A. m. ligustica* bees originated in Northern Italy (Fig. 1).

Here we ask whether and in how far selection for traits important for commercial beekeeping and natural adaptation to local climate have shaped sensory and cognitive capacities of the different honeybee subspecies.

## Materials and Methods

### Bees and hive management

For all of our honeybee subspecies we used the taxonomy of Engels (1999). Experiments were conducted in the summer of 2018 at the University of Würzburg (Bavaria, Germany). The experimental bees were derived from 23 queen-right colonies of six different subspecies of the Western honeybee *(Apis mellifera*). The colonies were headed by open mated queens from *A. m. carnica, A. m. ligustica, A. m. macedonica, A. m. mellifera, A. m. ruttneri* or *A. m. iberiensis* (n = 3-4 resp.). The queens were derived from populations central to the current distribution area of their respective subspecies (Fig. 1) and introduced into shook swarms of *A. m. carnica* at least ten weeks prior to data collection. All queens were shipped according to valid veterinary legislation and registered in the TRACES-Database. In order to prevent genetic pollution of the local honeybee population, all hives were equipped with excluder grids at the entrance, allowing only worker bees to fly freely. In addition, all drone brood was removed from the colonies, and the tip of one wing of each queen was clipped following the protocol of Human et al. (2015). In order to avoid drifting of homecoming forager bees between subspecies, the hive stands were grouped together according to the respective subspecies. For the same reason, within subspecies, only one pair of two differently colored hives with opposite flying directions were placed on each hive stand (Fig. 2). Despite the occurrence of some drought periods, the foraging conditions were comparatively good throughout the experimental period, with rich floral nectar flows and honey dew.

**Figure 2.**
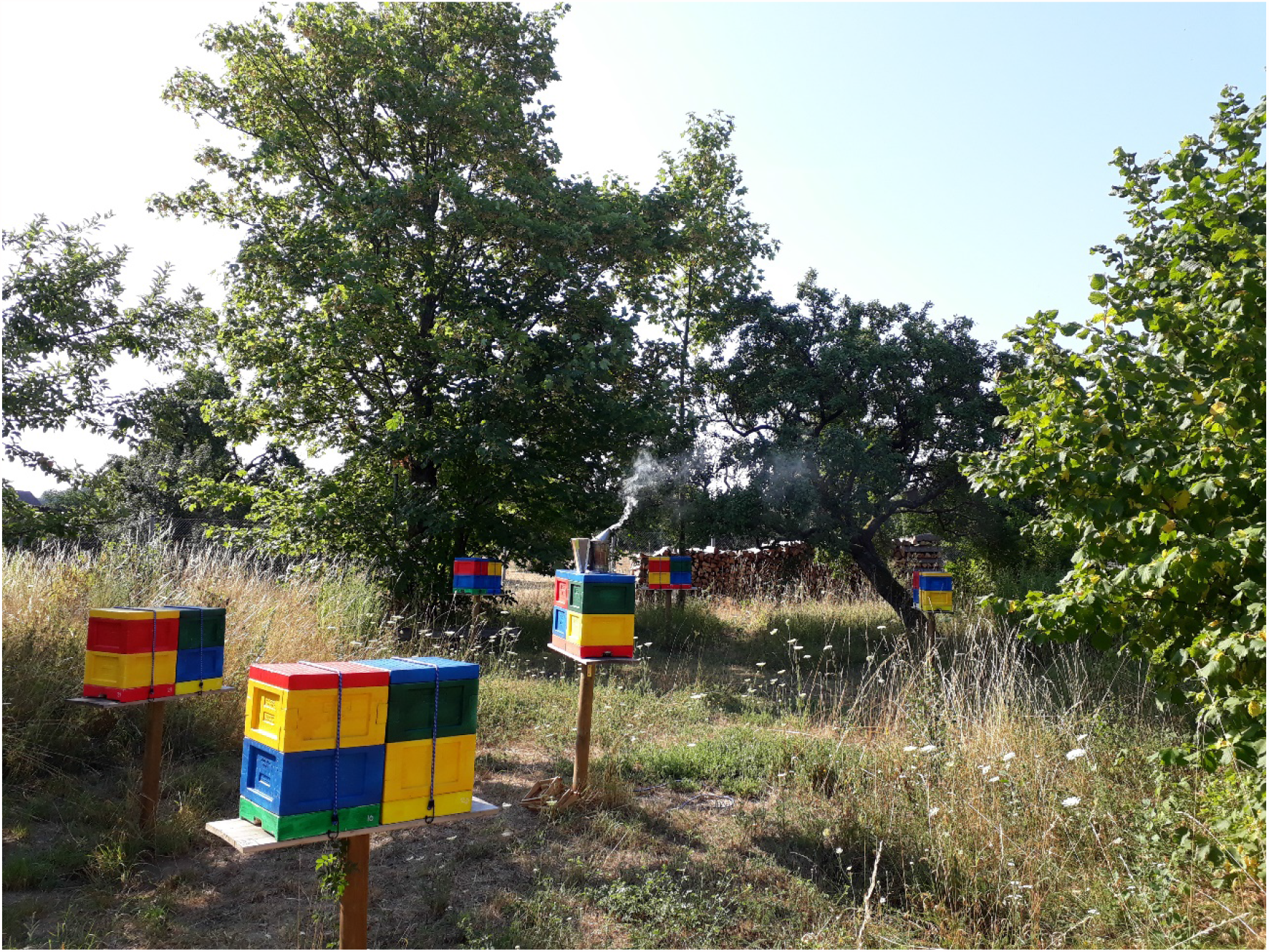
Location of experimental hives in a common apiary with four colonies of five subspecies of the Western honeybee *Apis mellifera* located in Southern Germany.

### Harnessing of bees

Returning foragers were collected at the hive entrance. Only foragers without pollen in their corbiculae were selected, assuming that they were nectar foragers. All bees were caught between morning and midday. Bees were caught individually in small glass vials, they were immobilized on ice and mounted in small brass tubes according to the standard protocol of Scheiner et al. (2013). Mounted bees were fed with 5 μl of a 30 % sucrose solution (according to Matsumoto et al., 2012) and placed in a dark incubator (temperature 35 °C, relative humidity 70 %) for one hour to recover from anesthesia and for sufficient motivation to learn (Scheiner et al. 2003b).

### Sucrose responsiveness

Sucrose responsiveness was tested prior to training as described elsewhere (Scheiner et al., 2001, 2003a, 2013, 2017; Hesselbach and Scheiner 2018). Briefly, both antennae of each bee were sequentially stimulated with water and a series of sucrose concentrations (0.1 %, 0.3 %, 1.0 %, 3.0 %, 10 %, 30 % w/v) in ascending order. The inter-trial interval was two min to prevent intrinsic sensitization (Scheiner et al. 2003b). It was recorded which sucrose concentration elicited the proboscis extension response (PER) for each bee. The total number of proboscis extension responses is the gustatory response score (GRS) of a bee (Scheiner et al., 2013).

### Appetitive olfactory learning and memory tests

To quantify associative olfactory learning and memory we used the protocol described in Scheiner et al. (2013). Only bees displaying no spontaneous response to the conditioned odor were tested. Briefly, bees were trained with 30 % sucrose solution as unconditioned stimulus and reward and 5 µl of 1-hexanol as conditioned stimulus (Sigma Aldrich, Steinheim, Germany). During each conditioning trial, we recorded which bee had already learned the conditioned stimulus and displayed proboscis extension before her antennae were touched with sucrose solution. The total number of conditioned proboscis extension responses of a bee constitutes her acquisition or learning score (Hesselbach and Scheiner 2018, Scheiner et al. 2013). The following number of bees were conditioned: n_*carnica*_ : 86, n_iberiensis_ : 56, n_*mellifera*_ : 71; n_*ruttneri*_ : 74; n_*macedonica*_ : 45; n_*ligustica*_ : 18.

### Statistics

All statistical analyses were conducted using SPSS Statistics 26 (IBM, Armonk, NY, USA). The GRS and acquisition scores were tested for normal distribution using the Shapiro-Wilk test. Since both scores were not distributed normally within each subspecies, non-parametric Kruskal Wallis H tests were performed with post hoc tests using p values corrected for multiple comparisons. To test for effects of colony on acquisition scores we also performed Kruskal Wallis H tests within each subspecies. The number of bees showing the correct response during the acquisition phase (i.e. acquisition curves) was analyzed using generalized linear models (logit function) using the binary responses in each acquisition trial as dependent variable and subspecies and GRS as factors. Correlations between GRS and acquisition scores were performed using Spearman rank correlation.

## Results

### Learning performance

Gustatory response scores (GRS) of bees trained to 1-hexanol were overall very high (Fig. 3A) and did not differ between subspecies (P > 0.05, Kruskal Wallis H Test), which indicated a high responsiveness to sucrose. Based on earlier experiments with *A*.*m. carnica* and *A*.*m. ligustica* we therefore expected high acquisition scores in all subspecies. We pooled data from foragers from different hives within each subspecies, because colony did not have an effect on acquisition score within each subspecies (P > 0.05, Kruskal Wallis H test).

**Figure 3.**
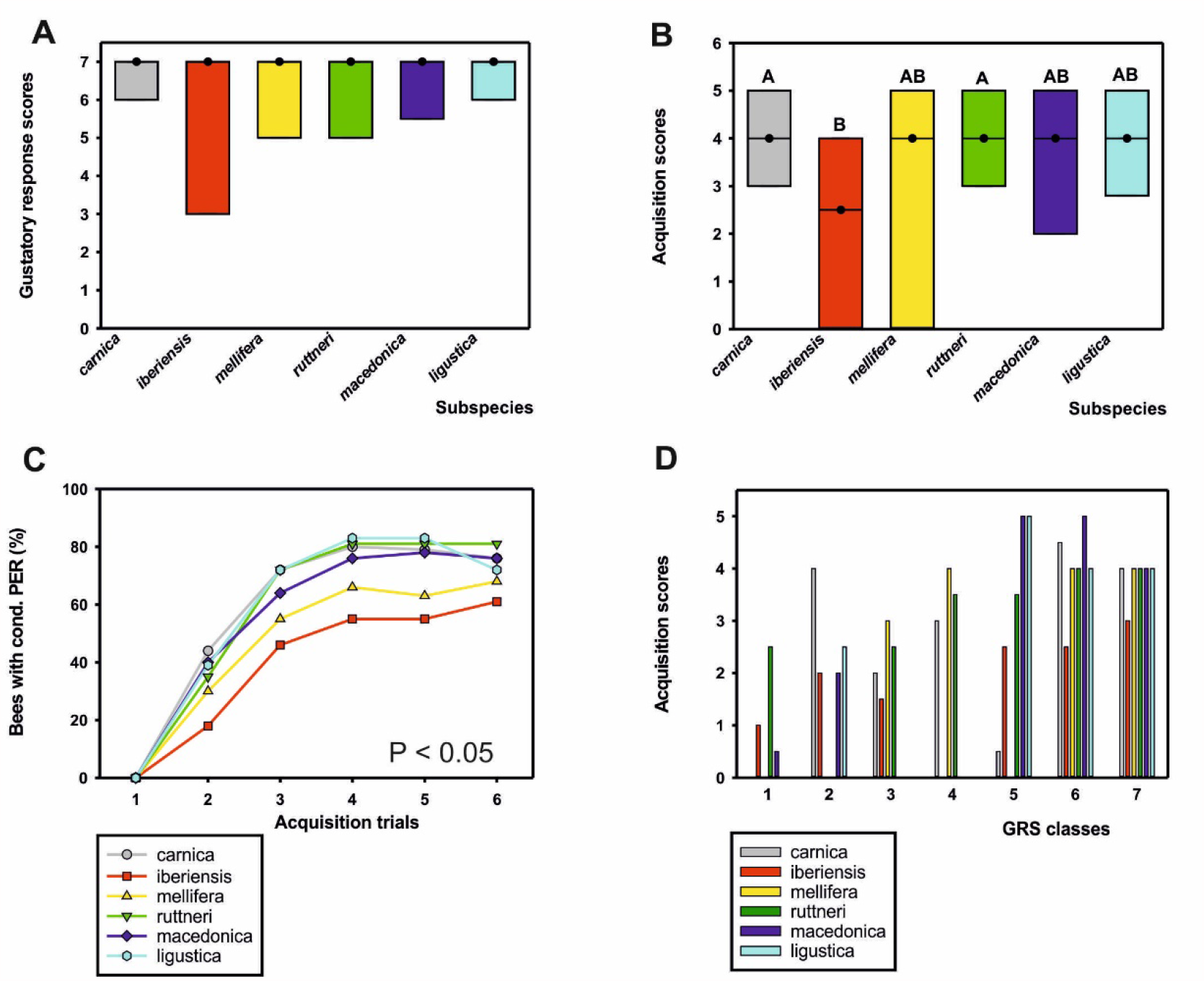
**A:** Sucrose responsiveness measured as gustatory response scores (GRS) of foragers belonging to the five different honeybee subspecies trained in this study. Median GRS (dots) and quartiles (25%: lower line, 75%: upper line) are presented. Subspecies were all highly responsive to sucrose and did not differ in their GRS (P > 0.05). **B**: Median acquisition scores (dots) of bees from subspecies trained in classical olfactory conditioning and quartiles (see A). Subspecies which differed significantly have a different letter. **C:** Acquisition curves of subspecies based on individual responses to the conditioned odor in each of the 6 training trials. There was a significant effect of subspecies on learning performance (P < 0.05). **D**. Correlation between GRS and learning scores in each subspecies. GRS were grouped on the x-axis. Median acquisition scores are shown for bees in each GRS class. Different subspecies are indicated by colors. Generally, the higher the GRS, the higher was the acquisition score, demonstrating a better learning performance. Number of bees tested see Table S1.

Comparison of acquisition scores yielded a significant effect of subspecies on learning performance (Fig. 3B; P < 0.01, Kruskal Wallis H Test). Foragers of the *A*.*m. iberiensis* subspecies displayed significantly lower acquisition scores than those of the *A*.*m. carnica* and *A*.*m. ruttneri* subspecies (P < 0.01 in each comparison). For a more detailed analysis of learning behavior, we compared the learning curves of the different subspecies and included GRS as within-subject factor in the model. Subspecies had a significant effect on learning curves (Fig. 3C; Chi^2^ _(5,349)_ = 12.75, P < 0.05, GLM). *A*.*m. iberiensis* foragers performed significantly more poorly than *A*.*m. carnica* (P < 0.05,) and *A*.*m. ruttneri* foragers (P < 0.05), while all of the other subspecies did not differ in their learning curve. GRS had a large and significant effect on learning curves (Chi^2^_(6,349)_ = 36.87; P < 0.001). Bees with higher GRS performed better than bees with lower GRS across subspecies. We further tested for a correlation between GRS and acquisition scores within each subspecies.

GRS correlated with acquisition scores significantly positively in the subspecies *A*.*m. carnica* (rho = 0.34, P < 0.001), *A*.*m. mellifera* (rho = 0.40, P < 0.001) and *A*.*m. ruttneri* (rho = 0.32, P < 0.01) but not in the other subspecies (Fig. 3D; P > 0.05, Spearman rank correlations). However, the same trend is observable in the other subspecies and it appears that the unequal distribution of GRS and partially a low sample size is related to the absence of a significant correlation.

## Discussion

This is the first study comparing the cognitive abilities of six different *Apis mellifera* subspecies from across Europe under standardized conditions in a common apiary. The bees tested for their appetitive olfactory learning performance were all highly motivated, i.e. displaying a similar and high sucrose responsiveness, which is an indicator of their “learning motivation” and can predict learning performance (Scheiner et al. 1999, 2001a, b, 2005, Scheiner, 2012). If the correlation between GRS and acquisition scores frequently demonstrated in *A*.*m. carnica* (Scheiner et al. 1999, 2005) and *A*.*m. ligustica* (Scheiner et al. 2001a,b) were also present in the other honeybee subspecies, there should be no or only little difference in the learning performance of the different subspecies. In fact, most of the European honeybee subspecies we analyzed did not differ in their learning performance from each other, supporting this hypothesis. However, *A*.*m. iberiensis*, a subspecies native to Southern Europe, performed significantly more poorly in our classical olfactory learning paradigm compared to *A*.*m. carnica* and *A*.*m. ruttneri* and displayed a non-significant tendency to perform less well than the other subspecies. This led us to question the general nature of the correlation between GRS and acquisition scores. We found a significant and expected positive correlation in *A*.*m. carnica, A*.*m. mellifera* and *A*.*m. ruttneri*. The reason why we did not see this correlation in *A*.*m. ligustica* might be due to the lower sample size in this experiment, but we nevertheless decided to include those data for comparison, and earlier experiments demonstrated a significant positive correlation in between GRS and acquisition scores in this subspecies, too (Scheiner et al. 2001a,b). In fact, the trend for a positive correlation between GRS and acquisition scores is observable in most subspecies tested (Fig. 3D), but the distribution of GRS was sometime suboptimal for correlation analyses, because most bees were highly responsive. Here, further experiments with larger samples sizes within each subspecies can help to ultimately show this correlation in all subspecies presented here.

Learning differences between different strains of honeybees (*A. m. ligustica*) selected for high or low amounts of stored pollen (Page and Fondrk, 1995), for example, could be explained by a different sucrose responsiveness, which correlated with the probability to collect pollen or nectar (Scheiner et al., 2001a,b). However, sucrose responsiveness did not differ between subspecies in this experiment, and we only tested nectar foragers (Fig. 3A). Thus we can exclude the possibility that *A*.*m. iberiensis* foragers were simply “less motivated” to learn compared to the *A*.*m. carnica* or *A*.*m. ruttneri* foragers. Therefore, the lower learning performance of this subspecies might be related to other factors including genetic differences related to foraging behavior. The differences in the learning performance of *A*.*m. iberiensis* foragers and foragers of the other subspecies appears not to be linked to the lineage of this subspecies, because *A*.*m. mellifera*, which is the second representative of lineage M, performs as well as the other subspecies.

It is conceivable that a differential adaptation to temperature-related stress may be linked to a different learning performance. Iberian bees might have a higher energy demand to cope with heat stress and / or higher foraging activity compared to honeybees from Central Europe, similar to what has been suggested by Iqbal et al. (Iqbal et al., 2019) for Arabian honeybees. A greater foraging activity, in turn, leads to accumulation of oxidative stress (Margotta et al., 2018) and reduces learning performance (Behrends et al., 2007; Scheiner and Amdam, 2009). However, *A*.*m. ruttneri* faces similar climatic stress because of high temperatures and still performed very well and significantly better than *A*.*m. iberiensis*. So, it is more likely that a combination of factors contributes to the lower learning performance of *A*.*m. iberiensis* compared to *A*.*m. carnica*.

Differential foraging strategies and related learning performance might be such a further reason for a differential performance of *A*.*m. carnica* and *A*.*m. iberiensis*. Diverse mechanisms involving learning appear to be important in decision making of individual foragers (e.g. Ferguson et al., 2001). Pérez-Claudio et al. (2018) discussed the lower learning ability of *A. m. syriaca* compared to *A*.*m. caucasia* in reverse association task in light of a higher predation rate in their native habitat. This might favor a risk-minimizing foraging strategy with a higher floral fidelity. Conversely, *A. m. caucasia* showed a higher flexibility in reverse association learning task, consistent with a previously described lower floral fidelity (Cakmak et al., 2010) and a lower predation rate in their foraging range (Pérez-Claudio et al., 2018). Whether the foraging strategy of *A*.*m. carnica* and the other subspecies differ from that of *A*.*m. iberiensis* is currently being investigated in our lab.

In an olfactory PER learning experiment comparing the performance of the Africanized honeybees (*Apis mellifera scutella*ta hybrid) with that of the Western honeybee *A. m. carnica*, the former performed significantly less well (Couvillon et al., 2010). A hypothesis which was posed by the authors was that the African honeybee might have been ecologically more successful than the Western honeybee, which might have been related to their lower learning performance. The authors suggest that if learning did not induce additional costs, there should be universal selection for high learning performance (Couvillon et al., 2010). However, a high degree of variation is maintained in natural populations of insects (McGuire and Hirsch, 1977), suggesting that learning could theoretically incur a fitness cost (Johnston, 1982; Dukas, 1999; Laughlin, 2001). Experiments in the fruit fly *Drosophila melanogaster* demonstrate an evolutionary trade-off between learning ability and competitive ability, supporting the hypothesis that selection for improved learning is consistently linked with a decreased competitive ability for limited food resources in larvae (Mery and Kawecki, 2003). A similar trade-off between improved learning performance and successful, aggressive strategies are conceivable for the honeybee. Africanized honeybees are notorious for their aggressive behavior. Similarly, the Iberian honeybee is typically more aggressive than the Carniolan honeybee (Adam, 1983; Ruttner, 1988), which was also apparent in our apiary, where all of the subspecies were hosted under equal climatic conditions.

A recent study comparing the learning performance of *A. m. carnica, A*.*m. ligustica* and *A*.*m. jemenitica* showed that the former two did not differ in their PER learning performance, similar to our findings, whereas *A. m. jemenitica* performed less well (Iqbal et al., 2019). In their experiments, the smaller body size of *A. m. jemenitica* was considered a possible reason for poorer learning, based on studies of body size and learning performance in bumble bees (Worden et al., 2005). However, it is debatable whether brain size is a good indicator of learning behavior, because there are different outcomes of experiments trying to link this factor with behavioral repertoires and cognitive abilities in animals (Couvillon et al., 2010; Chittka and Skorupski, 2011; Kotrschal et al., 2013). Further, it would not explain our differences between *A*.*m. iberiensis* and *A*.*m. carnica*, since they have a similar size (Ruttner, 1988). In addition, *A*.*m. ruttneri* is slightly smaller than either *A*.*m. carnica* or *A*.*m. iberiensis*. If size did matter, we would expect learning differences here, too. An intriguing hypothesis which awaits further investigation is that *A*.*m. iberiensis* bees differ from *A*.*m. carnica* bees and the other subspecies tested in their amount of neurotransmitters in the brain. A higher baseline brain titer of the biogenic amine octopamine, which itself is involved in the mediation of the reward to appetitive PER learning (Hammer and Menzel, 1998), might be responsible for a better learning performance. In support of this hypothesis, we could show that bees performing different social tasks (i.e. nurse bees vs. foragers or pollen collectors vs. nectar collectors), do not only differ in their titers of octopamine and its metabolic precursor tyramine (Reim and Scheiner, 2014; Scheiner et al., 2017a,b) but also in their appetitive learning capability (Scheiner et al. 1999; Scheiner et al., 2017a,b). Appetitive learning performance of honeybees can be improved by treating the bees with octopamine (Behrends and Scheiner, 2012) or tyramine (Scheiner et al., 2017b). Differential octopamine brain titers may be related to different foraging strategies of the different races and also to their aggression. This interesting neuroecological question awaits further study. Further, *A*.*m. iberiensis* foragers may have a reduced size of the mushroom bodies, important brain centers involved in learning and multimodal processing, or may have fewer synapses in their calyces of the mushroom bodies, leading to a reduced appetitive learning performance. This question has to be studied in future experiments.

## Conclusions

The most significant finding of our study is that differences in cognitive abilities are part of the intraspecific diversity in *A. mellifera*, similar to what has been demonstrated for other behavioral traits (Brillet et al., 2002; Kamel et al., 2003; Köppler et al., 2007; Uzunov, 2015a,b). While most of the subspecies from across Europe tested were very similar in associative learning capacities and the correlation between sucrose responsiveness and appetitive learning performance, the Iberian honeybee surprised with a reduced learning performance which was independent of the main motivational factor sucrose responsiveness. It could be linked to different foraging strategies, the higher aggressiveness of the Iberian bees, different amounts of neurotransmitters in the brain, a different size of brain neuropils important for learning or to genetic differences, all of which might have been shaped by ecological factors. It is an exciting open question how the neuroecology of foraging behavior and learning might thus be interlinked and shaped by adaptation to local climate and habitat.

## Acknowledgements

We thank our colleagues Dirk Ahrens, Cecilia Costa, Fani Hatjina, Dylan Elen, Thomas Galea, Alice Pinto and Léon Quiévy who provided queens of the different subspecies as well as their invaluable knowledge of the relevant biology. We also gratefully acknowledge the collaboration with Aleksandar Uzunov, who provided important comments on the study set up. Furthermore, we would like to thank Stefanie Schmidt and Wiebke Schäfer for help in data collection and comments on an earlier draft of the manuscript.

---

**Table S1.** Number of bees in each GRS class of each subspecies.

| GRS class | carnica | iberiensis | mellifera | ruttneri | macedonica | ligustica |
| --- | --- | --- | --- | --- | --- | --- |
| 1 | 3 | 5 | 5 | 2 | 4 | 0 |
| 2 | 4 | 5 | 7 | 2 | 3 | 2 |
| 3 | 6 | 6 | 3 | 6 | 1 | 0 |
| 4 | 3 | 0 | 2 | 4 | 0 | 0 |
| 5 | 2 | 4 | 3 | 6 | 3 | 1 |
| 6 | 8 | 4 | 14 | 3 | 4 | 5 |
| 7 | 60 | 32 | 37 | 51 | 30 | 10 |

## References

Adam, B. (1983). In search of the best strains of bees and the results of the evaluations of the crosses and races. Hamilton, Ill., U.S.A, Hebden Bridge, West Yorkshire, U.K: Dadant; Northern Bee Books.

Behrends, A., Scheiner, R. (2009). Evidence for associative learning in newly emerged honey bees (Apis mellifera). Animal Cognition, 12(2), 249–255. https://doi.org/10.1007/s10071-008-0187-7

Behrends, A., Scheiner, R. (2012). Octopamine improves learning in newly emerged bees but not in old foragers. Journal of Experimental Biology, 215(7), 1076–1083. https://doi.org/10.1242/jeb.063297

Behrends, A., Scheiner, R., Baker, N., Amdam, G. V. (2007). Cognitive aging is linked to social role in honey bees (Apis mellifera). Experimental Gerontology, 42(12), 1146–1153. https://doi.org/10.1016/j.exger.2007.09.003

Bouga, M., Alaux, C., Bienkowska, M., Büchler, R., Carreck, N. L., Cauia, E. et al. (2011). A review of methods for discrimination of honey bee populations as applied to European beekeeping. Journal of Apicultural Research, 50(1), 51–84. https://doi.org/10.3896/IBRA.1.50.1.06

Brillet, C., Robinson, G. E., Bues, R., Le Conte, Y. (2002). Racial differences in division of labor in colonies of the honey bee (Apis mellifera). Ethology, 108(2), 115–126. https://doi.org/10.1046/j.1439-0310.2002.00760.x

Cakmak, I., Song, D. S., Mixson, T. A., Serrano, E., Clement, M. L., Savitski, A., et al. (2010). Foraging response of Turkish honey bee subspecies to flower color choices and reward consistency. Journal of Insect Behavior, 23(2), 100–116. https://doi.org/10.1007/s10905-009-9199-7

Chen, C., Liu, Z., Pan, Q., Chen, X., Wang, H., Guo, H., et al. (2016). Genomic analyses reveal demographic history and tmperate adaptation of the newly discovered honey bee subspecies Apis mellifera sinisxinyuan n. Ssp. Molecular Biology and Evolution, 33(5), 1337–1348. https://doi.org/10.1093/molbev/msw017

Chittka, L., Skorupski, P. (2011). Information processing in miniature brains. Proceedings. Biological Sciences, 278(1707), 885–888. https://doi.org/10.1098/rspb.2010.2699

Couvillon, M. J., DeGrandi-Hoffman, G., Gronenberg, W. (2010). Africanized honeybees are slower learners than their European counterparts. Naturwissenschaften, 97(2), 153–160. https://doi.org/10.1007/s00114-009-0621-y

De La Rúa, .P, Galián, J., Serrano, J., Moritz, R. F. A. (2001). Genetic structure and distinctness of Apis mellifera L. populations from the Canary Islands. Molecular Ecology, 10(7), 1733–1742.

De La Rúa, P., Jaffé, R., Dall’Olio, R., Muñoz, I., Serrano, J. (2009). Biodiversity, conservation and current threats to European honeybees. Apidologie, 40(3), 263–284. https://doi.org/10.1051/apido/2009027

Dukas, R. (1999). Costs of memory: Ideas and predictions. Journal of Theoretical Biology, 197(1), 41–50. https://doi.org/10.1006/jtbi.1998.0856

Engel, M.S. (1999). The taxonomy of recent and fossil honey bees (Hymenoptera: Apidae; Apis). J Hymenopt Res, (8), 165–196.

Ferguson, HJ., Cobey S., Smith BH. 2001. Sensitivity to a change in reward is heritable in the honeybee, Apis mellifera. Animal Behaviour 61:527–534. DOI: 10.1006/anbe.2000.1635.

Hammer, M., Menzel, R. (1998). Multiple sites of associative odor learning as revealed by local brain microinjections of octopamine in honeybees. Learning & Memory, 5(1), 146–156.

Han, F., Wallberg, A., Webster, M. T. (2012). From where did the Western honeybee (Apis mellifera) originate? Ecology and Evolution, 2(8), 1949–1957.

Hesselbach H, Scheiner R. 2018. Effects of the novel pesticide flupyradifurone (Sivanto) on honeybee taste and cognition, Scientific Reports 8: 4954.

Human, H., Brodschneider, R., Dietemann, V., Dively, G., Ellis, J. D., Forsgren, E., et al. (2015). Miscellaneous standard methods for Apis mellifera research. Journal of Apicultural Research, 52(4), 1–53. https://doi.org/10.3896/IBRA.1.52.4.10

Iqbal, J., Ali, H., Owayss, A. A., Raweh, H. S. A., Engel, M. S., Alqarni, A. S., Smith, B. H. (2019). Olfactory associative behavioral differences in three honey bee Apis mellifera L. Races under the arid zone ecosystem of central Saudi Arabia. Saudi Journal of Biological Sciences, 26(3), 563–568. https://doi.org/10.1016/j.sjbs.2018.08.002

Jensen, A. B., Palmer, K. A., Boomsma, J. J., Pedersen, B. V. (2005). Varying degrees of Apis mellifera ligustica introgression in protected populations of the black honeybee, Apis mellifera mellifera, in northwest Europe. Molecular Ecology, 14(1), 93–106. https://doi.org/10.1111/j.1365-294X.2004.02399.x

Johnston, T. D. (1982). Selective costs and benefits in the evolution of learning. In Advances in the Study of Behavior (Vol. 12, pp. 65–106). Elsevier. https://doi.org/10.1016/S0065-3454(08)60046-7

Kamel, S. M., Strange, J. P., Sheppard, W. S. (2003). A scientific note on hygienic behavior in Apis mellifera lamarckii and A. m. carnica in Egypt. Apidologie, 34(2), 189–190. https://doi.org/10.1051/apido:2003014

Köppler, K., Vorwohl, G., Koeniger, N. (2007). Comparison of pollen spectra collected by four different subspecies of the honey bee Apis mellifera. Apidologie, 38(4), 341–353. https://doi.org/10.1051/apido:2007020

Kotrschal, A., Rogell, B., Bundsen, A., Svensson, B., Zajitschek, S., Brännström, I., et al. (2013). Artificial selection on relative brain size in the guppy reveals costs and benefits of evolving a larger brain. Current Biology, 23(2), 168–171. https://doi.org/10.1016/j.cub.2012.11.058

Laughlin, S. (2001). Energy as a constraint on the coding and processing of sensory information. Current Opinion in Neurobiology, 11(4), 475–480. https://doi.org/10.1016/s0959-4388(00)00237-3

Margotta, J. W., Roberts, S. P., Elekonich, M. M. (2018). Effects of flight activity and age on oxidative damage in the honey bee, Apis mellifera. Journal of Experimental Biology, 221(14).

Matsumoto, Y., Menzel, R., Sandoz, J.-C., Giurfa, M. (2012). Revisiting olfactory classical conditioning of the proboscis extension response in honey bees: A step toward standardized procedures. Journal of Neuroscience Methods, 211(1), 159–167. https://doi.org/10.1016/j.jneumeth.2012.08.018

McGuire, T. R., Hirsch, J. (1977). Behavior-genetic analysis of Phormia regina: Conditioning, reliable individual differences, and selection. Proceedings of the National Academy of Sciences of the United States of America, 74(11), 5193–5197. https://doi.org/10.1073/pnas.74.11.5193

Meixner, M. D., Costa, C., Kryger, P., Hatjina, F., Bouga, M., Ivanova, E., Büchler, R. (2010). Conserving diversity and vitality for honey bee breeding. Journal of Apicultural Research, 49(1), 85–92. https://doi.org/10.3896/IBRA.1.49.1.12

Meixner, M. D., Worobik, M., Wilde, J., Fuchs, S., Koeniger, N. (2007). Apis mellifera mellifera in eastern Europe – morphometric variation and determination of its range limits. Apidologie, 38(2), 191–197. https://doi.org/10.1051/apido:2006068

Mery, F., Kawecki, T. J. (2003). A fitness cost of learning ability in Drosophila melanogaster. Proceedings of the Royal Society B 270(1532), 2465–2469. https://doi.org/10.1098/rspb.2003.2548

Moritz, R. F. A., Kraus, F. B., Kryger, P., Crewe, R. M. (2007). The size of wild honeybee populations (Apis mellifera) and its implications for the conservation of honeybees. Journal of Insect Conservation, 11(4), 391–397. https://doi.org/10.1007/s10841-006-9054-5

Page, R. E., Fondrk, M. K. (1995). The effects of colony-level selection on the social organization of honey bee (Apis mellifera L.) colonies: Colony-level components of pollen hoarding. Behavioral Ecology and Sociobiology, 36(2), 135–144. https://doi.org/10.1007/BF00170718

Pérez-Claudio, E., Rodriguez-Cruz, Y., Arslan, O. C., Giray, T., Agosto Rivera, J. L., Kence, M., et al., (2018). Appetitive reversal learning differences of two honey bee subspecies with different foraging behaviors. PeerJ, 6, e5918. https://doi.org/10.7717/peerj.5918

Radloff, S. E., Hepburn, H. R., Hepburn, C., De La Rúa, P. (2001). Morphometric affinities and population structure of honey bees of the Balearic Islands (Spain). Journal of Apicultural Research, 40(3-4), 97–103.

Reim, T., Scheiner, R. (2014). Division of labour in honey bees: age-and task-related changes in the expression of octopamine receptor genes. Insect Molecular Biology, 23(6), 833–841. https://doi.org/10.1111/imb.12130

Ruottinen, L., Berg, P., Kantanen, J., Kristensen, T. N., Praebel, A. (2014). Status and conservation of the Nordic brown bee. NordGen, 42.

Ruttner, F. (1988). Biogeography and taxonomy of honeybees. Berlin, Heidelberg: Springer-Verlag.

Ruttner, F., Tassencourt, L., J. Louveaux (1978). Biometrical-statistical analysis of the geographic variability of Apis mellifera L. Apidologie, 9(4), 363–381. https://doi.org/10.1007/s003590050360

Scheiner, R. (2012). Birth weight and sucrose responsiveness predict cognitive skills of honeybee foragers. Animal Behaviour, 84(2), 305–308. https://doi.org/10.1016/j.anbehav.2012.05.011

Scheiner, R., Amdam, G. V. (2009). Impaired tactile learning is related to social role in honeybees. Journal of Experimental Biology 212(7), 994–1002. https://doi.org/10.1242/jeb.021188

Scheiner, R., Erber, J. (2009). Sensory thresholds, learning and the division of foraging labor in the honey bee. Organization of Insect Societies: From Genomes to Socio-Complexity. Harvard University Press, Cambridge, 335–356.

Scheiner, R., Abramson, C. I., Brodschneider, R., Crailsheim, K., Farina, W. M., Fuchs, S., et al. (2013). Standard methods for behavioural studies of Apis mellifera. Journal of Apicultural Research, 52(4), 1–58. https://doi.org/10.3896/IBRA.1.52.4.04

Scheiner, R., Barnert, M., Erber, J. (2003a). Variation in water and sucrose responsiveness during the foraging season affects proboscis extension learning in honey bees. Apidologie, 34(1), 67–72. https://doi.org/10.1051/apido:2002050

Scheiner, R., Entler, B. V., Barron, A. B., Scholl, C., Thamm, M. (2017). The effects of fat body tyramine level on gustatory responsiveness of honeybees (Apis mellifera) differ between behavioral castes. Frontiers Systems Neuroscience 11.https://doi.org/10.3389/fnsys.2017.00055

Scheiner, R., Erber, J., Page, R. E. (1999). Tactile learning and the individual evaluation of the reward in honey bees (Apis mellifera L.). Journal of Comparative Physiology A, 185(1), 1–10.

Scheiner, R., Kuritz-Kaiser, A., Menzel, R., Erber, J. (2005). Sensory responsiveness and the effects of equal subjective rewards on tactile learning and memory of honeybees. Learning & Memory, 12(6), 626–635. https://doi.org/10.1101/lm.98105

Scheiner R, Müller U, Heimburger S, Erber J. (2003b). Activity of protein kinase A and gustatory responsiveness in the honey bee (Apis mellifera L.). Journal of Comparative Physiology A 189: 427–434.

Scheiner, R., Page, R. E., Erber, J. (2001a). Responsiveness to sucrose affects tactile and olfactory learning in preforaging honey bees of two genetic strains. Behavioural Brain Research, 120(1), 67–73. https://doi.org/10.1016/S0166-4328(00)00359-4

Scheiner, R., Page, R.E., Erber, J. (2001b). The effects of genotype, foraging role and sucrose perception on the tactile learning performance of honey bees (Apis mellifera L.). Neurobiology of Learning and Memory 76, 138–150.https://doi.org/10.1006/nlme.2000.3996

Scheiner, R., Page, R. E., Erber, J. (2004). Sucrose responsiveness and behavioral plasticity in honey bees (Apis mellifera). Apidologie, 35(2), 133–142. https://doi.org/10.1051/apido:2004001

Scheiner, R., Reim, T., Sovik, E., Entler, B. V., Barron, A. B., Thamm, M. (2017). Learning, gustatory responsiveness and tyramine differences across nurse and forager honeybees. Journal of Experimental Biology, 220(8), 1443–1450. https://doi.org/10.1242/jeb.152496

Sheppard, W. S., Arias, M. C., Grech, A., Meixner, M. D. (1997). Apis mellifera ruttneri, a new honey bee subspecies from Malta. Apidologie, 28(5), 287–293. https://doi.org/10.1051/apido:19970505

Sheppard, W. S., Meixner, M. D. (2003). Apis mellifera pomonella a new honey bee subspecies from Central Asia. Apidologie, 34(4), 367–375. https://doi.org/10.1051/apido:2003037

Uzunov, A., Costa, C., Panasiuk, B., Meixner, M., Kryger, P., Hatjina, F., et al. (2015a). Swarming, defensive and hygienic behaviour in honey bee colonies of different genetic origin in a pan-European experiment. Journal of Apicultural Research, 53(2), 248–260. https://doi.org/10.3896/IBRA.1.53.2.06

Uzunov, A., Meixner, M. D., Kiprijanovska, H., Andonov, S., Gregorc, A., Ivanova, E. et al., (2015b). Genetic structure of Apis mellifera macedonica in the Balkan Peninsula based on microsatellite DNA polymorphism. Journal of Apicultural Research, 53(2), 288–295. https://doi.org/10.3896/IBRA.1.53.2.10

van Engelsdorp, D., & Meixner, M. D. (2010). A historical review of managed honey bee populations in Europe and the United States and the factors that may affect them. Journal of Invertebrate Pathology, 103 Suppl 1, 95. https://doi.org/10.1016/j.jip.2009.06.011

Worden, B. D., Skemp, A. K., Papaj, D. R. (2005). Learning in two contexts: The effects of interference and body size in bumblebees. Journal of Experimental Biology, 208(11), 2045–2053. https://doi.org/10.1242/jeb.01582

Zammit-Mangion, M., Meixner, M., Mifsud, D., Sammut, S., Camilleri, L. (2017). Thorough morphological and genetic evidence confirm the existence of the endemic honey bee of the Maltese Islands Apis mellifera ruttneri: Recommendations for conservation. Journal of Apicultural Research, 56(5), 514–522. https://doi.org/10.1080/00218839.2017.1371522

